# Design and assessment of TRAP-CSP fusion antigens as effective malaria vaccines

**DOI:** 10.1101/613653

**Authors:** Chafen Lu, Gaojie Song, Kristin Beale, Jiabin Yan, Emma Garst, Emily Lund, Flaminia Catteruccia, Timothy A. Springer

## Abstract

The circumsporozoite protein (CSP) and thrombospondin-related adhesion protein (TRAP) are major targets for pre-erythrocytic malaria vaccine development. However, the most advanced CSP-based vaccine RTS,S provides only partial protection, highlighting the need for innovative approaches for vaccine design and development. Here we design and characterize TRAP-CSP fusion antigens, and evaluate their immunogenicity and protection against malaria infection. TRAP N-terminal folded domains were fused to CSP C-terminal fragments consisting of the C-terminal αTSR domain with or without the intervening repeat region. Homogenous, monomeric and properly folded fusion proteins were purified from mammalian transfectants. Notably, fusion improved expression of chimeras relative to the TRAP or CSP components alone. Immunization of BALB/c mice with the *P. berghei* TRAP-CSP fusion antigens formulated in AddaVax adjuvant elicited antigen-specific antibody responses. Remarkably, fusion antigens containing the CSP repeat region conferred complete sterile protection against *P. berghei* sporozoite challenge, and furthermore, mice that survived the challenge were completely protected from re-challenge 16 weeks after the first challenge. In contrast, fusion antigens lacking the CSP repeat region were less effective, indicating that the CSP repeat region provided enhanced protection, which correlated with higher antibody titers elicited by fusion antigens containing the CSP repeat region. In addition, we demonstrated that N-linked glycans had no significant effect on antibody elicitation or protection. Our results show that TRAP-CSP fusion antigens could be highly effective vaccine candidates. Our approach provides a platform for designing multi-antigen/multi-stage fusion antigens as next generation more effective malaria vaccines.

## Introduction

Malaria remains a global health problem with an estimated 216 million cases of infection and 445,000 deaths worldwide in 2016. Children under age 5 are most vulnerable to malaria infection and suffer high mortality. The most advanced vaccine, RTS,S, reduced infection by 27% in infants and 46% in children during the first 18 months, and protection declined thereafter [1, 2], leaving an unmet need for more effective vaccines.

Malaria is caused by *Plasmodium* parasites transmitted by *Anopheles* mosquitoes. Infected mosquitoes introduce salivary gland sporozoites into the host during a blood meal. Sporozoites infect hepatocytes, and subsequent infection of red blood cells causes the symptoms of malaria. Most vaccine development has targeted the pre-erythrocytic stage (liver stage infection). Both subunit vaccines and live sporozoites, attenuated by radiation or mutation, or given in combination with chemoprophylaxis, have been studied in preclinical and clinical trials. Although immunization with live sporozoites provides high levels of protection [3–6], the cost of manufacturing sporozoites from infected mosquitos, maintaining them viably, and the requirement for multiple intravenous injections have prevented wide applications in malaria endemic regions.

Two major proteins on the surface of sporozoites, the circumsporozoite protein (CSP) and thrombospondin-related adhesion protein (TRAP), are the focus of pre-erythrocytic subunit vaccine development; both are essential for sporozoite motility and liver-stage infection [7, 8]. CSP, the most abundant surface protein in sporozoites, is composed of an N-terminal domain (NTD), Region I (RI), where CSP is cleaved during cell invasion [9, 10], a central repeat region, and the C-terminal αTSR domain followed by a glycosylphosphatidylinositol (GPI) membrane anchor (Fig 1A). The RTS vaccine includes the C-terminal portion of the repeat region (R) and the αTSR domain (T) from *P. falciparum* CSP (Fig 1A) fused to the hepatitis B surface antigen (S). TRAP consists of the N-terminal von Willebrand factor type-1 (VWA) or integrin I-domain, the thrombospondin type-1 repeat (TSR) domain, C-terminal repeats, and transmembrane and cytoplasmic domains (Fig 1A). TRAP delivered by adenovirus prime and modified vaccinia Ankara virus (MVA) boost regimens achieved 40-95% protection efficacy in murine models [11, 12], and 21% sterile protection in one phase I/II trial [13], and in another clinical trial reduced the risk of infection by 67% [14]. With the limited protective efficacy of the RTS,S vaccine and vaccines targeting TRAP in clinical trials, combination vaccination targeting both CSP and TRAP has recently been explored. Although a phase I/II clinical trial combining adjuvanted TRAP and RTS,S showed no benefit of protection from TRAP [15], a recent preclinical study showed that combining a modified RTS,S-like vaccine R21 with viral vectored TRAP significantly enhanced protective efficacy compared to single component vaccines [16]. However, immunological interference has been a concern when combining two vaccines. Bauza et al. reported that co-administration of TRAP and CSP vaccines in one immunization regimen resulted in a reduction of CSP antibodies and no improvement of protection over either subunit vaccine alone [12]. In another study, CD8+ T cell interference was observed when combining viral vectored CSP with the blood stage antigen merozoite surface protein 1 (MSP1) [17]. Further work is needed to tailor and refine immunization regimens to achieve the additive benefit of CSP and TRAP co-vaccination.

**Figure 1.**
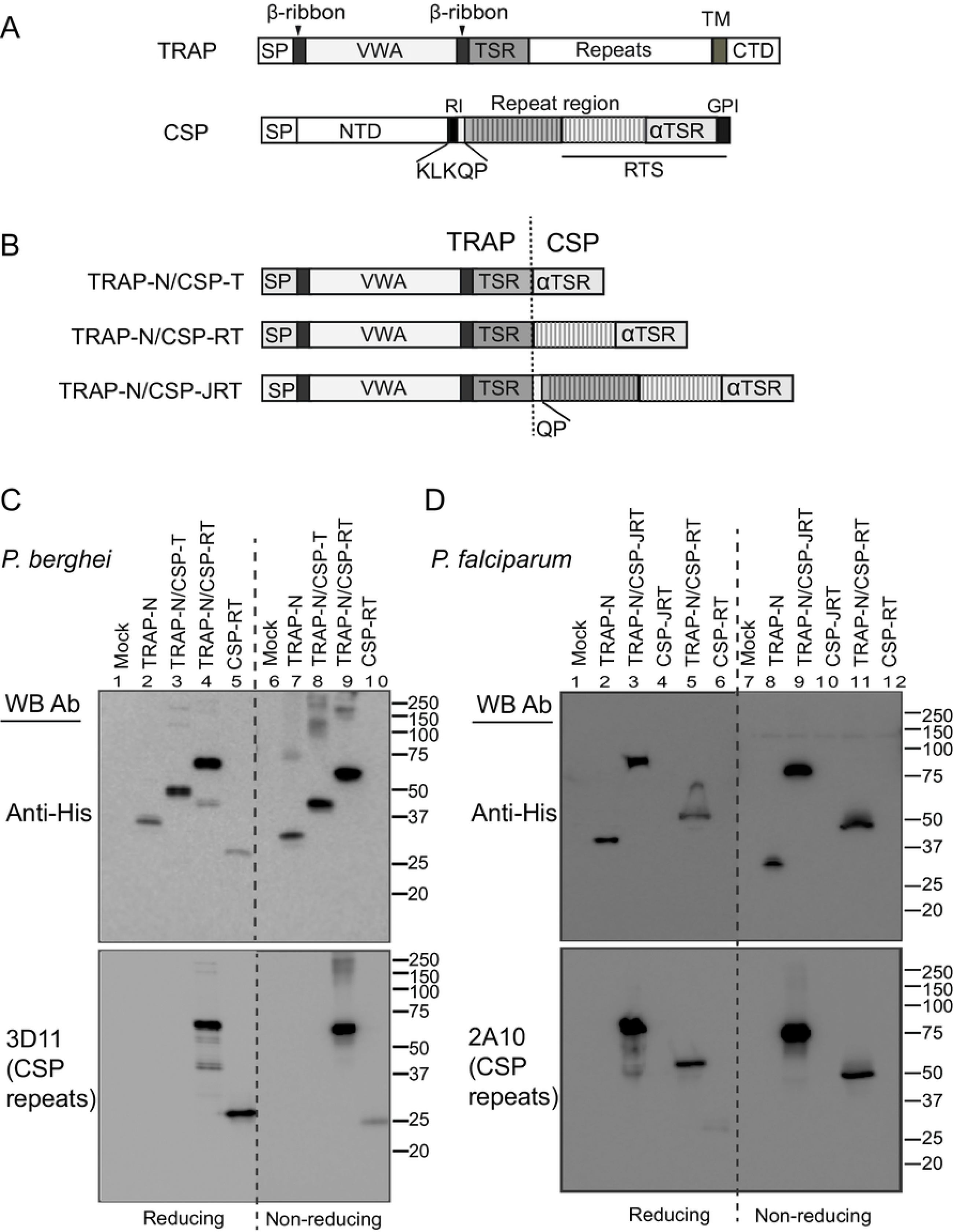
TRAP-CSP fusion antigen design and expression. (A and B) Schematic diagrams. SP, signal peptide; TM, transmembrane domain; CTD, cytoplasmic domain; NTD, N-terminal domain; RI, region I; GPI, glycosylphosphatidylinositol membrane anchor. The region included in the RTS component of RTS,S vaccine is shown under the CSP diagram. The dashed line shows the fusion junction between TRAP and CSP. (C) and (D) Expression in 293T transfectants of *P. berghei* constructs (C) and *P. falciparum* constructs (D). Supernatants from 293T cells transiently transfected with the indicated constructs or vector alone (mock) were subjected to 10% reducing or nonreducing SDS-PAGE and Western blot using antibodies to the His tag or the CSP repeat region as indicated.

Here we aimed to design TRAP-CSP fusion antigens to provide the benefit of protection conferred by delivering both antigens as a single protein. TRAP-CSP chimeric constructs containing functional protein domains and regions with protective B and T cell epitopes of both proteins were generated, and monomeric and properly folded chimeric proteins were purified from mammalian transfectants. Immunization with TRAP-CSP fusion antigens elicited strong antigen-specific antibody responses and sterile immunity against *P. berghei* challenge in mice. Our results show that TRAP-CSP fusion antigens offer promise as more effective vaccine candidates.

## Materials and methods

### DNA constructs

TRAP and CSP protein sequences are from *P. berghei* strain ANKA (GeneBank accession number AAB63302 for TRAP and CDS44911 for CSP) and from *P. falciparum 3D7* (accession number AAA29775 for TRAP and CAB38998 for CSP). Synthetic cDNAs were codon optimized for mammalian expression by Atum Bio (Newark, CA) for *P. berghei* constructs and by GenScript (Piscataway, NJ) for *P. falciparum* constructs. *P. berghei* chimeras and TRAP-N constructs contain TRAP amino acids Gln^25^- Pro^291^, with amino acids 99-119 (KRYGSTSKASLRFIIAQLQNN) replaced by the equivalent sequence from *P. falciparum* (HSDASKNKEKALIIIKSLLST, α2 and α3 helices in VWA domain structure) [18] due to unsuccessful expression of the native *Pb*TRAP sequence. *P. berghei CSP* fusion fragments contain CSP amino acids Asn^164^- Ser^318^ and Pro^239^-Ser^318^ for TRAP-N/CSP-RT and TRAP-N/CSP-T chimeras, respectively, with Asn^280^ mutated to Ser to remove the putative N-glycosylation site. *P. falciparum* chimeras contain TRAP amino acids Arg^26^-Asp^297^ with the nonconserved Cys^55^ and the N-glycosylation site Asn^132^ mutated to Ser and Gln, respectively, and CSP Gln^96^- Ser^375^ and Asn^207^- Ser^375^, for the TRAP-N/CSP-JRT and TRAP-N/CSP-RT, respectively. *P. berghei* constructs were inserted in pLexM vector [18]. *P. falciparum* constructs were cloned into Nhe I-Bam HI sites of the pIRES2-EGFP vector (Takara Bio, formerly Clontech). All constructs contain a modified murine kappa chain secretion signal peptide and C-terminal His tag [18], and were confirmed by DNA sequencing.

### Antibodies

Monoclonal antibodies to *P. falciparum* TRAP were generated and characterized as described in the Supplemental Methods and Figures. Antibodies 1G12, 4B3 and 4C2 to *P. falciparum* CSP were kindly provided by Dr. Nicholas J. MacDonald (NIH, Malaria Vaccine Development Branch). 3D11 and 2A10 hybridoma lines were obtained from BEI Resources (ATCC), and IgG antibodies were purified from serum-free culture supernatants using protein G affinity chromatography. Secondary antibodies were rabbit polyclonal anti-His (Delta Biolabs), Horseradish peroxidase (HRP)-anti-rabbit and HRP-anti-mouse IgG (GE Healthcare) for Western blot, and HRP-penta-His antibody (Qiagen) and HRP-anti-mouse IgG (Abcam) for ELISA.

### Cell culture

293T cells (ATCC) were cultured in DMEM medium supplemented with10% fetal bovine serum (FBS). 293S GnTI- cells (obtained from Philip J. Reeves Laboratory, Departments of Biology and Chemistry, Massachusetts Institute of Technology) [19] and Expi293F cells (Thermo Fisher Scientific) were cultured in suspension in serum-free Ex-Cell 293 medium (Sigma) and Expi293 medium (Thermo Fisher Scientific), respectively.

### Protein expression and purification

293T cells in 6-well tissue culture plates were transfected using Lipofectamine^2000^ according to manufacturer’s instruction (Thermo Fisher Scientific). For scaleup transient transfection of 293S GnTI- cells, suspension cultures were transfected using polyethyleneimine [20]. Culture supernatants were harvested 6 days later. For stable transfection of Expi293F cells, adherent Expi293F cells were transfected in DMEM medium with 10% FBS using Lipofectamine^2000^ transfection. Selection was started 48 hours later by addition of 0.5 mg/ml G418 (final concentration). After 10-12 days, cells were harvested and sorted for top 5% GFP positive cells on a FACSAria machine (BD Biosciences). Cells were sorted a 2^nd^ time to further enrich GFP expressing cells. Sorted cells were expanded in suspension culture in serum-free Expi293 medium for protein purification. Proteins were purified from culture supernatants of transfectants by Ni-NTA followed by gel filtration chromatography as described [18]. For endoglycosidase H (Endo H) treatment, Ni-NTA purified materials were buffer exchanged to 50 mM sodium acetate, pH 5.5, and 150 mM NaCl, and digested with Endo H at 1:20 mass ratio of enzyme:protein at 4°C, overnight. Fractions from gel filtration chromatography were subjected to non-reducing SDS-PAGE and monomeric protein peak fractions were pooled and stored in aliquots at −80°C.

### Western blot

Cultural supernatants (10 μl) from transiently transfected 293T cells were mixed with 2.5 μl 5x Laemmli sample buffer containing 25% β-mercaptoethanol or 25 mM N-ethylmaleimide for reducing and non-reducing SDS-polyacrylamide gel electrophoresis (PAGE), respectively. Blotting to PVDF membrane was carried out using Tran-Blot Turbo transfer system (Bio-RAD). Membrane was probed with 0.4 μg/ml primary antibody, followed by incubation with HRP-conjugated 2^nd^ antibodies and chemiluminescence imaging using LAS-4000 system (Fuji Film). ImageJ software was used for quantitation of protein bands.

### Enzyme-linked immunosorbent assay (ELISA)

96-well Elisa plates (Costar) were coated with 50 μl of purified antibodies at 5 μg/ml in 50 mM sodium carbonate buffer, pH 9.5, 50 μl/well for 2 hrs at 37°C, and blocked with 3% BSA for 90 min at 37°C. His tagged TRAP or TRAP-CSP fusion proteins (50 μl, 0.4 μg/ml) was added and incubated at 4°C overnight. Binding was detected with HRP-anti-His (Penta-His Ab at1:5000 dilution) or with biotin-labeled primary antibody at 0.5 μg/ml followed by HRP-streptavidin. 10 min after addition of peroxidase substrate (Life Technologies), plates were read at 405 nM on an Emax plate reader (Molecular Devices).

### Immunization

10 μg *P. berghei* TRAP/CSP fusion antigens diluted in 100 μl PBS was mixed with 100 μl AddaVax adjuvant (InvivoGen, San Diego), and completely emulsified by pushing the mixture between two glass syringes via a two-way valve. BALB/c mice (The Jackson Laboratory) were injected intraperitoneally (i.p) with 200 μl of antigen and adjuvant emulsion per mouse. Mice were immunized 3 times (prime + 2 boosts) at intervals of three weeks. Control mice received PBS and adjuvant emulsion. Tail blood (50-100μl) was collected from each mouse for measuring antibody responses.

### Antibody titer measurement

Serum was obtained from tail blood and stored at −80°C until use. 96-well ELISA plates were coated with 50μl of TRAP-N/CSP-RT or TRAP-N/CSP-T antigen at 2.5 μg/ml in sodium carbonate buffer, pH 9.5, overnight at 4°C. Plates were blocked with 3% BSA. Sera were serially diluted 5-fold starting from 1:200, and 50 μl/well added in duplicate and incubated for 2 hrs at room temperature. Similarly, positive control antibody 3D11 to the *P. berghei CSP* repeat region or antibody to the His tag at the C-terminus of the antigens were used at 1 μg/ml. Serum from adjuvant alone immunized mice was diluted and added to each plate as negative control. After incubation with HRP-anti-mouse whole IgG, peroxidase substrate was added, and absorbance at 405 nm was read 10 min later. A semi-logarithmic dilution curve (x-axis: log dilution and y-axis: OD_405_, subtracted by the negative control) was generated for each serum sample, and a line parallel to the x axis was drawn at half of the OD value of the 3D11 or His tag positive control antibodies. Antibody titer was taken as the dilution factor where the dilution curve intercepted with the line at half of the OD value of the positive control.

### *P. berghei-infected* mosquitoes

PbGFP_CON_, a recombinant *Plasmodium berghei* (ANKA strain) that constitutively expresses GFP was used [21]. To infect mosquitoes, PbGFP_CON_-infected mice (3-7% parasitemia) were anaesthetized and laid over a cage of female *Anopheles stephensi* mosquitoes (50-100 mosquitoes per cage), which were allowed to feed for 15 min. Successful mosquito infection was confirmed 10 days after blood feeding by dissecting the midgut and examining under a fluorescent microscope for the presence of oocyst. At day 20 after blood feeding, infected mosquitoes (prevalence of infection >80%) were used for challenge.

### Challenge by infected mosquito bite

For challenge infection by mosquito bite, immunized or naive mice were anaesthetized and exposed to the bites of PbGFP_CON_-infected mosquitoes for 15 min as described above. Mice were monitored for blood stage infection at days 7, 9 and 12 post challenge by Giemsa staining of thin blood smears and microscopic examination, and parasitemia was determined as % infected red blood cells. Mice that remained blood stage parasite-free after 12 days were considered sterilely protected.

### Ethics Statement

Animal work was conducted in accordance with and was approved by the Harvard Medical School Institutional Animal Care and Use Committee (IACUC) under protocol #05010. Animals were cared in compliance with the U.S. Department of Agriculture (USDA) Animal Welfare Act (AWA) and the Public Health Service (PHS) Policy on Humane Care and Use of Laboratory Animals.

## Results

### Design of TRAP and CSP fusion antigens

TRAP and CSP fusion constructs combined functional domains defined in crystal structures and regions containing protective B and T-cell epitopes from both proteins (Fig 1B). The folded VWA and TSR domains at the N-terminus of TRAP [18] and the αTSR domain at the C-terminus of CSP [22] were left intact in all fusion constructs, which varied in their content of CSP repeats in between the folded domains (Fig 1B). The VWA and TSR domains of TRAP are required for sporozoite motility and host cell invasion [7, 23]. Moreover, the VWA domain is the target of potent protective CD8+ T cell responses elicited by TRAP or whole sporozoite immunization [24–26]. The CSP repeat region contains immunodominant B-cell epitopes recognized by sporozoite neutralizing antibodies, whereas the αTSR domain contains several T-cell epitopes associated with protection [26–30]. The junction (J) between the NTD and the repeat region, which is included in the TRAP-N/CSP-JRT construct and not in the RTS,S vaccine, has recently been identified to contain epitopes for potent protective antibodies [31–33]. Fusion constructs from both *P. berghei* and *P. falciparum* were generated for evaluation of protein expression and folding. *P. berghei* fusion proteins contain 4 putative N-glycosylation sites in the TRAP fragment for comparison of glycosylated and N-glycan shaved versions, whereas the single N-glycosylation site in the *P. falciparum* chimeric constructs was removed by mutating TRAP Asn^132^ to Gln.

### Expression, purification and folding state of TRAP-CSP fusion proteins

Expression of fusion proteins or fragments of TRAP and CSP alone were tested in mammalian 293 cell transfectants, which were previously used to obtain well-folded, glycosylated *P. falciparum* and *P. vivax* TRAP N-terminal fragments and CSP C-terminal fragments for structure and carbohydrate determination [18, 22, 34]. Equal amounts of culture supernatants from transiently transfected 293 cells were subjected to Western blot using antibodies to the C-terminal His-tag or to the CSP repeat region. TRAP-N protein from both *P. berghei* and *P. falciparum* was detected by anti-His, and migrated faster under non-reducing than reducing conditions, as typical of proteins with disulfide bonds (Fig 1C, lanes 2 and 7 and 1D, lanes 2 and 8). Lower amount of *P. berghei* CSP-RT fragment was observed (Fig 1C, lanes 5 and 10), whereas *P. falciparum* CSP-JRT and CSP-RT fragments were undetectable by anti-His (Fig 1D, top, lane 4 and 6, and lanes 10 and 12). However, a weak *P. falciparum* CSP-RT band of ~27 kD was detected by repeat region antibody 2A10 under reducing condition (Fig 1D, bottom, lane 6). In contrast to TRAP and CSP fragments alone, larger amounts of TRAP-CSP fusion proteins of both *P. berghei* and *P*. *falciparum* were detected by the His tag antibody and had the expected sizes (Fig 1C top, lanes 8 and 9 vs 7; Fig 1D top, lane 9 and 11 vs 8); quantification of protein bands under non-reducing showed 5- and 3- fold increases of *Pb*TRAP-N/CSP-RT and *Pb*TRAP-N/CSP-T, respectively, relative to *Pb*TRAP-N protein, and 3- and 2-fold increases of *Pf*TRAP-N/CSP-JRT and *Pf*TRAP-N/CSP-RT, respectively, relative to *Pf*TRAP-N. Additionally, the chimeras containing the CSP repeat region were recognized by CSP repeat region antibodies (Fig 1C and 1D, bottom). Thus, fusion improved expression of the chimeric proteins relative to the TRAP or CSP components alone.

TRAP-CSP fusion proteins were purified for further characterization. *P. berghei* fusion constructs, TRAP-N/CSP-RT and TRAP-N/CSP-T, containing 4 putative N-glycosylation sites, were expressed in 293S GnTI- cells, which lack N-acetylglucosaminyltransferase I (GnTI) and produce glycoproteins with short and homogeneous high mannose-type N-linked glycans that can be removed by endoglycosidase H (Endo H) [19], and purified by Ni-NTA affinity chromatography and treated with Endo H. Endo H removed ~6 kD mass from each of the *P. berghei* TRAP-N/CSP-RT and TRAP-N/CSP-T proteins and reduced their mass to 52 kD and 41 kD, respectively (Fig 2A, reducing SDS-PAGE). A second purification step, gel filtration chromatography, was utilized to obtain homogenous and monomeric proteins. A representative chromatogram is shown in Fig 2B. The purified *P. berghei* TRAP-N/CSP-RT and TRAP-N/CSP-T chimeras and their respective N-glycan shaved versions (designated dNG) each showed a single band by non-reducing SDS-PAGE (Fig 2C), confirming homogeneity and monomeric state. The *P. falciparum* TRAP-N/CSP-JRT and TRAP-N/CSP-RT fusion proteins, with the single N-glycosylation site mutated, were purified from Expi293F stable transfectants, and monomeric proteins showed mass of ~75 kD and 58 kD, respectively, under reducing condition (Fig 2D).

**Figure 2.**
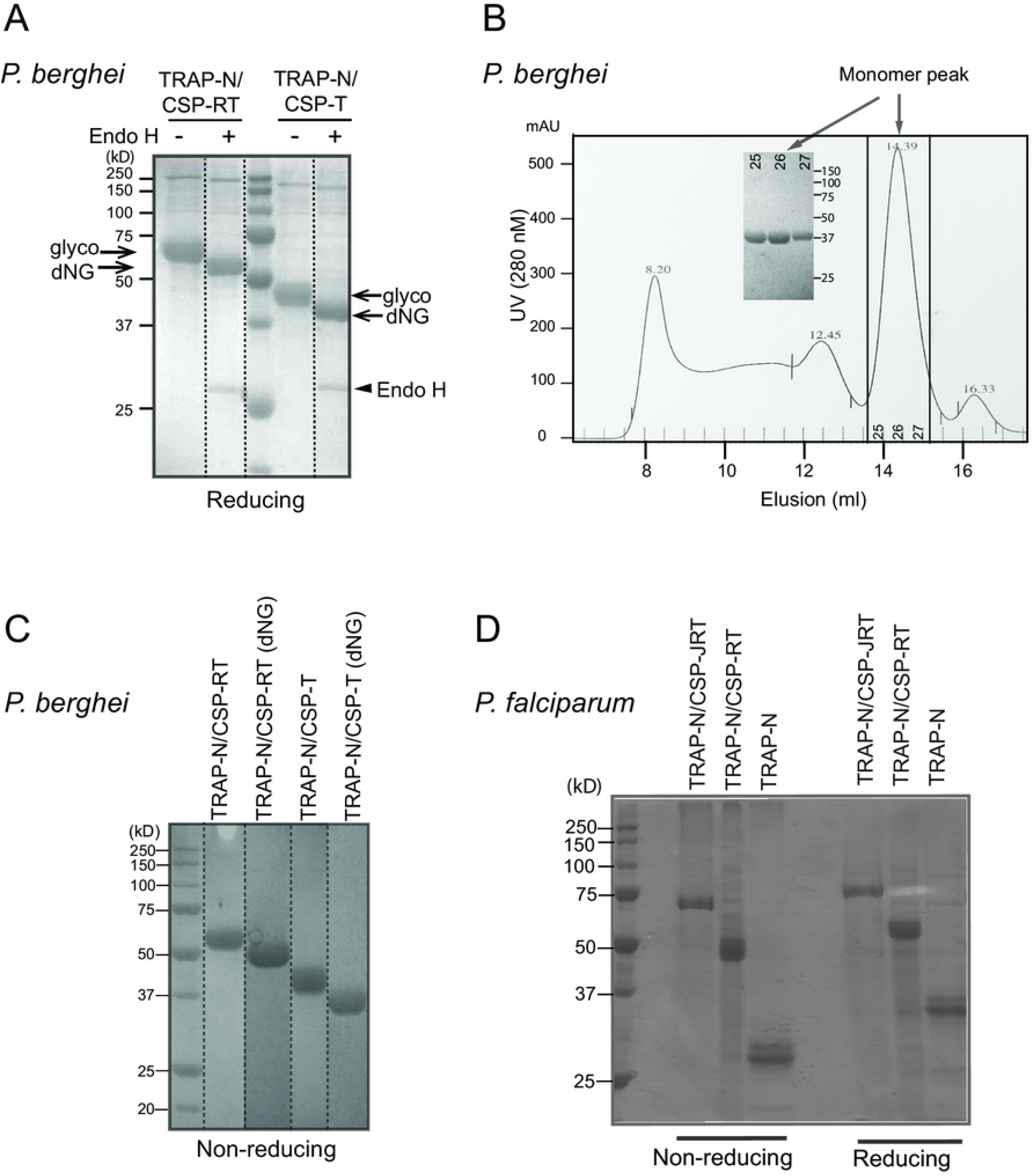
TRAP-CSP fusion protein purification. (A) *P. berghei* chimeras were produced by transient transfection of 293S GnTI^−^ cells, purified by Ni-NTA affinity chromatography, treated with or without Endo H, and subjected to reducing SDS-PAGE and Coomassie blue staining. Arrows point to glycosylated (glyco) and Endo H-treated (dNG) chimera and Endo H protein bands. Dotted lines separate lanes run on the same gel and moved. (B) Representative Superdex 200 10/300 GL column purification chromatogram, shown TRAP-N/CSP-T sample that had previously been purified by Ni-NTA and treated with Endo H. Insert, nonreducing SDS-PAGE of monomer peak fractions. (C) Nonreducing SDS-PAGE and Coomassie blue staining of *P. berghei* fusion proteins after the Superdex S200 purification step. (dNG) denotes Endo H treatment. Dotted lines divide lanes run on two identical gels and moved. (D) SDS-PAGE and Coomassie blue staining of purified *P. falciparum* TRAP-N/CSP-JRT and TRAP-N/CSP-RT fusion proteins run under nonreducing and reducing conditions.

Proper folding of recombinant antigens is required to elicit B cell responses to native proteins on parasites. Purification of monomeric well-behaved material already suggested that the fusion proteins were well folded. We utilized available, well-characterized monoclonal antibodies (mAbs) to *P. falciparum* TRAP and CSP to further probe the folding state of TRAP-CSP fusion proteins. We generated and characterized a panel of mAbs to *Pf*TRAP (Supplemental Material). CL2/1 and CL5/27 mAbs to PfTRAP VWA domain and CL5/22, CL5/42, CL5/46 and CL5/54 mAbs to the TSR domain and β-ribbon (Fig s1) recognized TRAP on the surface of transfectants (Fig s2, Table s1) and stained unfixed *P. falciparum* sporozoites (Fig s3, Table s1), suggesting that they recognize native epitopes. Furthermore, the epitopes of mAbs CL2/1, CL5/27, CL5/22, CL5/46 and CL5/54 are sensitive to disulfide reduction (Fig s5). The 4B3, 4C2 and 1G12 mAbs to the αTSR domain of *Pf*CSP have been shown to recognize CSP with intact disulfides and inhibit sporozoite invasion of liver cells *in vitro* [35]. The purified, monomeric *P. falciparum* TRAP-N/CSP-JRT and TRAP-N/CSP-RT fusion proteins bound to the VWA domain mAbs CL2/1 and CL5/27 and mAbs CL5/22, CL5/42, CL5/46 and CL5/54 to the TSR domain and β-ribbon at levels comparable to the TRAP-N alone (Fig 3A). Furthermore, the two fusion proteins reacted with the αTSR domain mAbs 4B3, 4C2 and 1G12 to a level comparable to the His tag antibody and the repeat region antibody 2A10 (Fig 3B). These results indicate that the TRAP VWA and TSR domains and the CSP αTSR domain of the fusion proteins are correctly folded.

**Figure 3.**
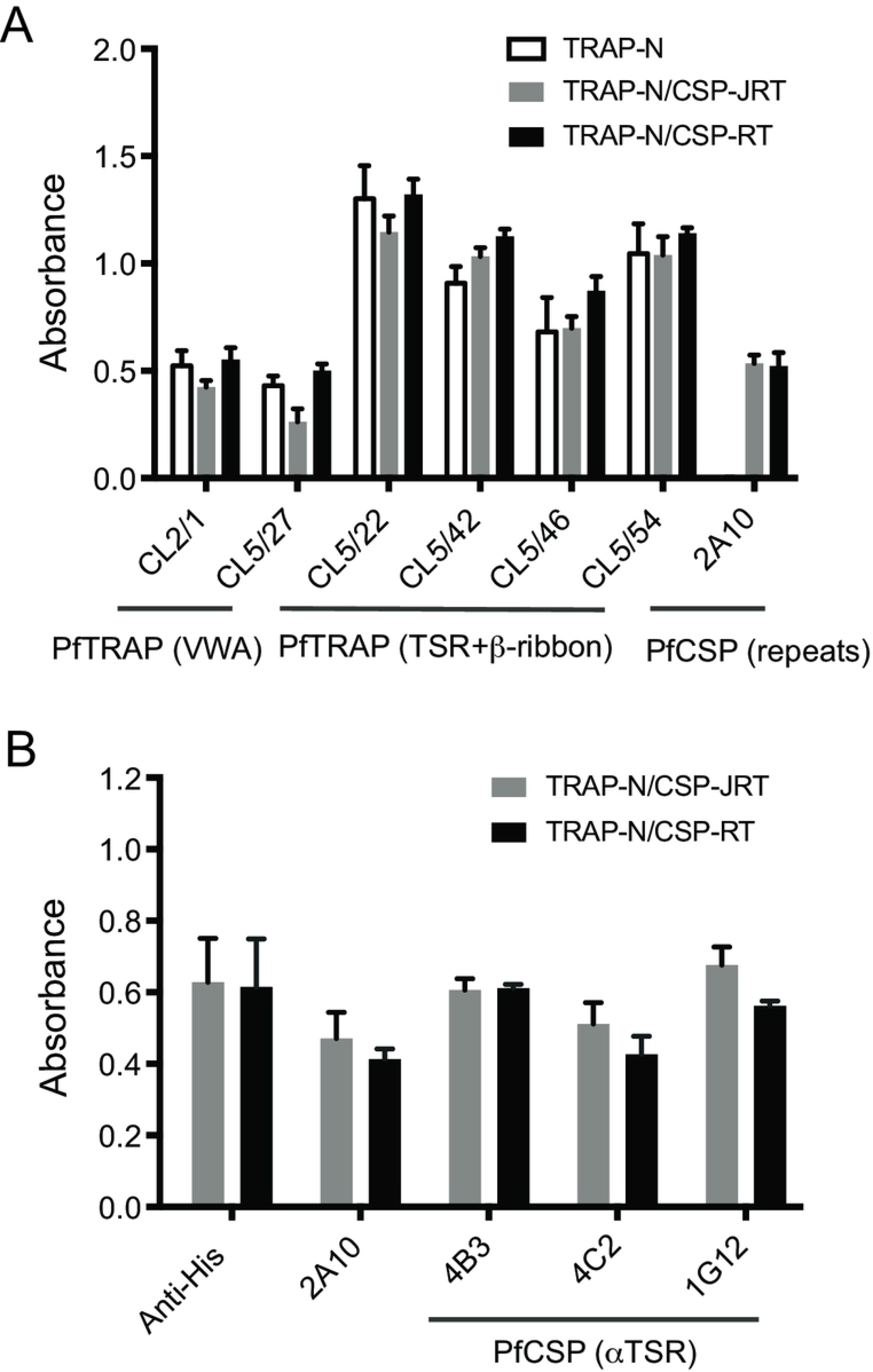
Reactivity of fusion proteins to antibodies that recognize correctly folded TRAP and CSP. ELISA plates were coated with the indicated antibodies and incubated with purified *P. falciparum* TRAP-CSP fusion proteins or TRAP-N alone as indicated. Binding to the immobilized antibodies was detected by HRP-penta-His antibody to the C-terminal His tag (A) or biotin-labeled TRAP antibody CL5/42 followed by HRP-streptavidin (B). Results are mean ± SD of triplicate wells, and representative of 3 independent experiments.

These results show that TRAP-CSP chimeras are well expressed in mammalian cells, are properly folded, and yielded homogenous, monomeric proteins after Ni-NTA and gel filtration chromatography.

### Humoral responses elicited by TRAP-CSP fusion antigens

Mice were immunized with *P. berghei* TRAP-N/CSP-T and TRAP-N/CSP-RT fusion antigens to evaluate immunogenicity and protection against *P. berghei* infection. To investigate the effect of the high mannose N-linked glycans on these chimeras, mice were also immunized with the Endo H-shaved versions, TRAP-N/CSP-RT (dNG) and TRAP-N/CSP-T (dNG), in which all but one N-acetylglucosamine residue was removed. Fusion antigens were formulated in AddaVax, a squalene-based oil-in-water emulsion. BALB/c mice were primed with10 μg adjuvanted antigen, followed by two boost immunizations. Sera were collected at weeks 6, 16 and 32 post antigen priming to determine antibody responses. Antigen-specific IgG antibody titers, defined as serum dilutions giving half of the absorbance of a positive control antibody, were measured by ELISA.

Whether the chimeras contained high-mannose or shaved N-glycans had little effect on antibody responses. Serum IgG antibody titers from the TRAP-N/CSP-RT immunized group were comparable to the TRAP-N/CSP-RT (dNG) antigen group, at all three time points (Fig 4A). Similarly, no significant differences in antibody titers were observed between the TRAP-N/CSP-T and TRAP-N/CSP-T (dNG) groups (Fig 4A and 4B). The results showed that the four putative high mannose type N-linked glycans in the TRAP-N fragment neither enhanced nor compromised antigen-specific antibody responses.

**Figure 4.**
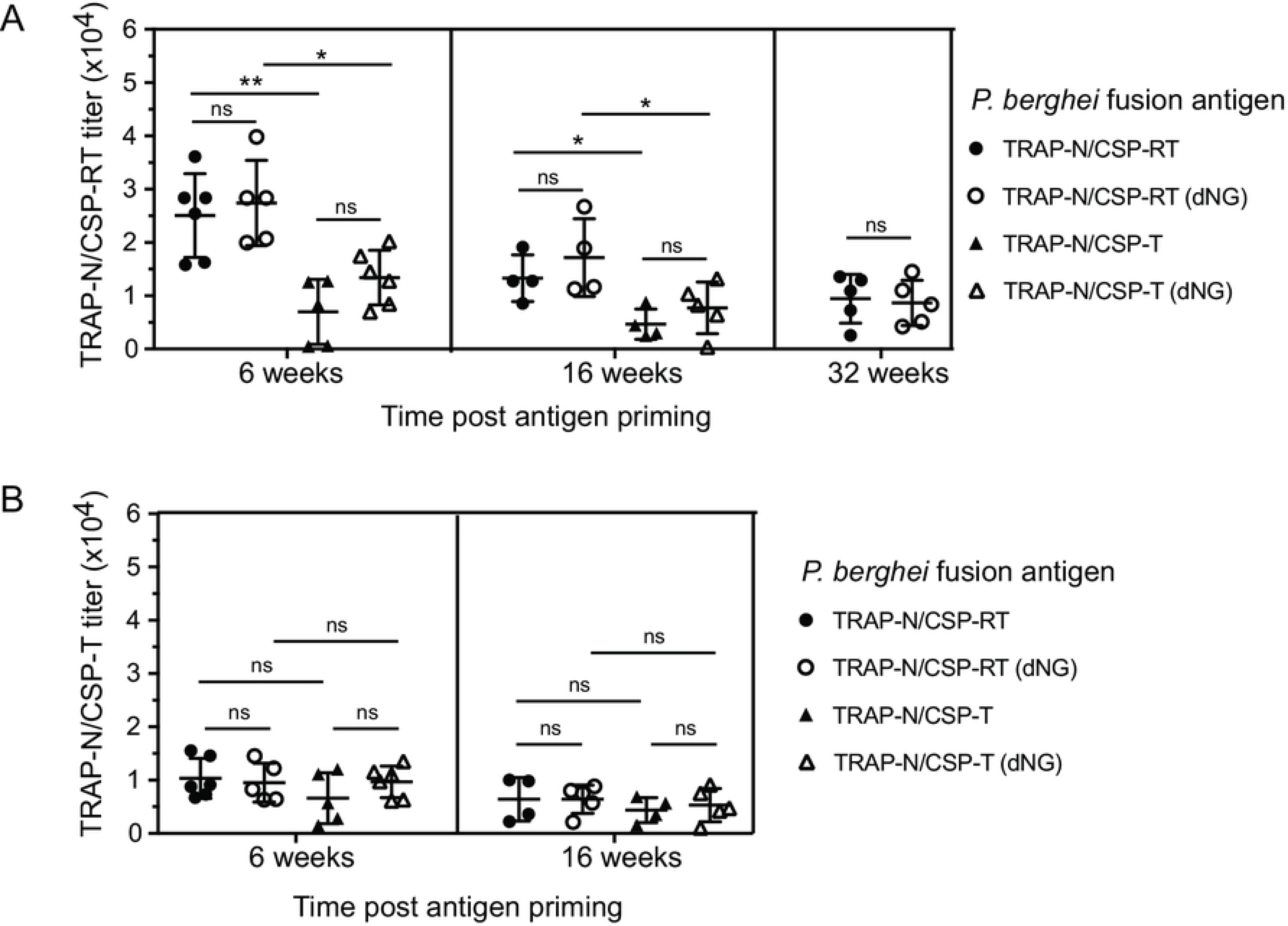
Antibody responses elicited by immunization with *P. berghei* TRAP-CSP fusion antigens. BALB/c mice were immunized with TRAP-N/CSP-RT and TRAP-N/CSP-T antigens and their respective N-glycan shaved versions (dNG) in AddaVax adjuvant. Tail blood was collected at the indicated time points post antigen priming and antigen specific IgG antibody titers were measured by ELISA. Antibody titers of individual mice were measured against the TRAP-N/CSP-RT antigen (A) and the TRAP-N/CSP-T antigen (B). Mean and SD are shown, and analyzed by One-way ANOVA with Sidak’s multiple comparisons test (GraphPad Prism 7). *, P<0.05; **, P<0.01; ns, not significant.

In contrast, the CSP repeats present in TRAP-N/CSP-RT compared to TRAP-N/CSP-T significantly enhanced antibody responses. As shown in Fig 4A, when antibody titers were measured against the TRAP-N/CSP-RT protein containing the repeat region, mice immunized with TRAP-N/CSP-RT and TRAP-N/CSP-RT (dNG) had ~2-fold higher titers than mice immunized with the TRAP-N/CSP-T and TRAP-N/CSP-T (dNG) antigens, 6 weeks and 16 weeks post priming (Fig 4A). By comparison, antibody titers against the TRAP-N/CSP-T protein lacking the repeat region were comparable among the four groups of mice (Fig 4B).

### Protection against *P. berghei* infection

Protection of immunized mice against *P. berghei* infection was assessed. Mice were challenged by bite with *P. berghei* infected *Anopheles stephensi* mosquitoes. Blood stage infection was monitored at days 7, 9 and 12 post challenge. Remarkably, 5 out of 5 mice immunized with TRAP-N/CSP-RT and TRAP-N/CSP-RT (dNG) antigens were completely free of blood stage parasites day 12 after challenge (100% sterile protection) (Table I). In contrast, the TRAP-N/CSP-T and TRAP-N/CSP-T (dNG) antigens each conferred 40% sterile protection; 3 of 5 challenged mice in each immunized group showed blood stage infection at each of days 7, 9, and 12 after challenge (Table I).

**Table I.** Protection against *P. berghei* challenge.

| *P. berghei*<br>fusion antigen<br>Immunization | 1<sup>st</sup> challenge<br>(16 weeks post antigen priming) |  |  |  | Re-challenge of protected mice<br>(32 weeks post antigen priming) |  |  |  |
| --- | --- | --- | --- | --- | --- | --- | --- | --- |
|  | Number<br>infected/<br>challenged<br>mice | Parasitemia (%)<sup>a</sup><br>(day post challenge) |  |  | Number<br>infected/<br>challenged<br>mice | Parasitemia (%)<sup>a</sup><br>(day post challenge) |  |  |
|  |  | Day 7 | Day 9 | Day 12 |  | Day 7 | Day 9 | Day 12 |
| TRAP-N/<br>CSP-RT | 0/5 | 0 | 0 | 0 | 0/5 | 0 | 0 | 0 |
| TRAP-N/<br>CSP-RT (dNG) | 0/5 | 0 | 0 | 0 | 0/5 | 0 | 0 | 0 |
| TRAP-N/<br>CSP-T | 3/5 | 3.5<br>2.9<br>2.3 | 3.8<br>4.7<br>3.9 | 8.4<br>10.4<br>8.5 | 2/2 | 2.4<br>3.1 | 3.8<br>4.6 | 9.1<br>10.5 |
| TRAP-N/<br>CSP-T (dNG) | 3/5 | 2.6<br>2.4<br>1.6 | 3.2<br>2.8<br>3.1 | 10.9<br>12.5<br>13.9 | 2/2 | 2.7<br>2.9 | 3.2<br>3.7 | 13.9<br>12.6 |
| None (naïve) | 1/1 | 3.1 | 8.0 | 18.9 | 1/1 | 2.3 | 3.8 | 13.8 |

Mice were immunized in AddaVax adjuvant and challenged as described in the Methods. Blood stage infection was monitored at days 7, 9 and 12 post each challenge. Mice free of blood stage infection 12 days after challenge were determined as sterilely protected. Mice sterilely protected from the 1<sup>st</sup> challenge were re-challenged 16 weeks after the 1<sup>st</sup> challenge.
<sup>a</sup> Data shown are individual infected mice; "0", no infection of the entire group.

To evaluate maintenance of protection, sterilely protected mice were re-challenged 16 weeks after the 1^st^ challenge (32 weeks after antigen priming). Strikingly, the TRAP-N/CSP-RT and TRAP-N/CSP-RT (dNG) immunized groups were completely protected from re-challenge (Table I). In contrast, the 2 mice each immunized with TRAP-N/CSP-T and TRAP-N/CSP-T (dNG) antigens that were protected at week 16 were not protected at week 32 and showed blood stage infection (Table I).

These results show that immunization with TRAP-N/CSP-RT and its N-glycan shaved (dNG) version conferred sustained sterile protection, whereas TRAP-N/CSP-T and its dNG version, lacking the CSP repeat region, were less protective. Thus, the CSP repeat region enhanced protection, which correlated with higher antibody titers elicited by the TRAP-N/CSP-RT and TRAP-N/CSP-RT (dNG) fusion antigens (Fig 4A).

## Discussion

The benefit of protection against malaria infection by co-immunization with CSP and TRAP was first demonstrated in 1991 by Khusmith *et al*., who showed that immunization of mice with a mixture of *P. yoeli* CSP and TRAP-expressing cells provided complete protection, whereas immunization with either cell type alone only provided partial protection [36]. Despite this result, pre-erythrocytic vaccine development has focused on CSP and TRAP single subunit vaccines over the last two decades and clinical trials to date with CSP and TRAP subunit vaccines have achieved only partial protection. We show here that combining TRAP and CSP fragments into a single fusion protein that contained well-folded protein domains and regions with protective T- and B-cell epitopes from each fusion partner yielded a vaccine that stimulated sterilizing immunity.

The TRAP-N/CSP-RT fusion antigen containing from *Pb*TRAP the VWA and TSR domains and from *Pb*CSP the C-terminal half of the repeat region and the αTSR domain conferred durable and complete sterile protection against *P. berghei* infection in BALB/c mice, a significant improvement over the 50% sterile protection obtained in BALB/c mice with three immunizations of adjuvanted full-length CSP protein [37]. By comparison, the *Pb*TRAP-N/CSP-T antigen that lacked the CSP repeat region was less protective (40% sterile protection), showing that the repeat region provided the enhanced protection, which correlated with higher antibody responses elicited by the TRAP-N/CSP-RT antigen. The same results on sterilizing immunity and antibody responses were obtained with chimeras containing high mannose or Endo H-shaved N-glycans. It is well known that the CSP repeats elicit strong sporozoite neutralizing antibody responses; several potent protective antibodies have been isolated and characterized [38–41]. On the other hand, TRAP delivered by viral vectors primarily stimulates CD8+ cytotoxic T cell responses that eliminate infected hepatocytes [11, 13, 42]. Further studies are required to understand how the VWA and TSR domains of TRAP and the repeats and αTSR domain of CSP additively or synergistically contribute to the sterile protection conferred by the TRAP-N/CSP-RT fusion antigen.

We expressed the TRAP-CSP fusion proteins in a mammalian system that allows proper disulfide formation and glycosylation. For protein-based subunit vaccines, proper folding of recombinant antigens to elicit antibody responses to native protein epitopes is important, and correct disulfide formation to stabilize antigen native conformation is critical. Many targets of malaria vaccine candidates, including TRAP and CSP, contain multiple disulfide bonds, which pose challenges to obtain antigens with correct disulfide formation from bacterial expression. Yeast and insect cell expression systems have achieved some success [35, 43, 44]. However, RTS,S is expressed in the yeast cytoplasm where disulfide bond formation is not promoted, and RTS has not been demonstrated to be monomeric or to have a natively folded CSP αTSR domain [45–48]. We showed that mammalian expressed, purified *P. falciparum* TRAP-CSP fusion proteins are properly folded by demonstrating their reactivity to a panel of antibodies that recognize correctly folded VWA and TSR domains of *Pf*TRAP and the αTSR domain of *Pf*CSP. We expect the purified monomeric *P. berghei* TRAP-CSP fusion proteins are properly folded likewise.

Importantly, mammalian cells mannosylate and fucosylate TSR domains similarly to *Plasmodium* sporozoites. X-ray crystallography and mass spectrometry have shown that the TSR domains of *Plasmodium* TRAP or its orthologue in *Toxoplasma gondii*, when expressed in mammalian cells, bear a C-linked mannose on tryptophan and O-linked fucose on threonine [18, 49]. Both modifications are also found on TRAP on the surface of *P. falciparum* sporozoites [34]. Furthermore, the αTSR domain of CSP expressed in mammalian cells is fucosylated [22], as is the αTSR domain of intact CSP on the surface of *P. falciparum* sporozoites [34]. Yeast lacks the gene required for fucosylation of TSR domains and the αTSR domain expressed in *Pichia* was not fucosylated [22]. Fucosylation and mannosylation of TSR domains in Apicomplexans have only recently been recognized [22] [18, 34, 49] and are little considered if at all in the malaria vaccine literature. However, these modifications may be of considerable importance in stabilizing folding and enhancing production of components of subunit vaccines and also may constitute important portions of both B cell folded antigenic epitopes and T cell peptide epitopes.

Less is known about N-glycosylation in *Plasmodium*. In contrast to N-glycans in yeast, insect and mammalian cells, *Plasmodium* makes severely truncated N-glycans composed of one or two N-acetylglucosamine (GlcNAc) residues [50]. Glycans can shield native epitopes from immune responses, as shown by the impact of the HIV-1-glycan shield on antibody elicitation [51, 52]. Efforts to overcome glycosylation differences between native and recombinant malarial antigens have been mostly restricted to mutations to remove putative N-glycosylation sites [43, 44]. We compared immune responses in mice to *P. berghei* TRAP-N/CSP-RT and TRAP-N/CSP-T fusion proteins with high mannose N-glycans to chimeras with all but one N-acetylglucosamine residue removed by Endo H (dNG). We found comparable antibody responses and no effect on protection in our *P. berghei* challenge model. Datta *et al* reported that N-glycosylation of the *Pf*s25 antigen delivered as DNA vaccine did not significantly affect antibody response and malaria transmission blocking efficacy [53]. In contrast, un-glycosylated *P. falciparum* MSP1 antigen derived from transgenic mice and glycosylated high mannose MSP1 from baculovirus expression both protected monkeys from *P. falciparum* challenge, whereas glycosylated MSP1 antigen from mice conferred no protection [54]. These results suggest that the types of N-glycans, i.e. complex type with terminal sialic acids from mammalian cells vs. high mannose type from insect cells, contributed to the difference in protection. Therefore, the impact of N-glycosylation of malarial antigens on protection efficacy may vary both among antigens and the types of glycans.

Given the highly encouraging protection efficacy conferred by the *P. berghei* TRAP-N/CSP-RT fusion antigens in this study, TRAP-CSP chimeras offer promise as effective preerythrocytic vaccines and as fusion partners for multistage chimeric vaccines. Recently identified potent protective and “dual specificity” antibodies to *Pf* CSP bind to the NANP repeats and the junction (J) region [32, 33]. This region has not been included in our *P. berghei* constructs or RTS,S. However, the *P. falciparum* chimeric construct containing this region, TRAP-N/CSP-JRT, was expressed even better than the TRAP-N/CSP-RT construct (Fig 1D). It would be very interesting to compare protection efficacy of TRAP-N/CSP-JRT with the TRAP-N/CSP-RT chimera in a *P. falciparum* challenge model. A chimeric antigen of *P. vivax* CSP and the transmission-blocking vaccine target s25 conferred 43% protection against parasite infection and 82% transmission blocking efficacy [55]. Recent crystal structures of HAP2, a gamete fusion protein that is also a transmission blocking target, offer further promise for rational design of TRAP-CSP-HAP2 fusion antigens with dual efficacy [56–59].

In conclusion, we have shown that TRAP-CSP fusion antigens could be highly effective vaccine candidates. Chimeric antigens provide a platform for development of multi-antigen/multi-stage vaccines to elicit dual immunity to prevent infection of humans and block transmission of infection by mosquitos using a single immunization regimen.

## Acknowledgments

We thank Dr. Shahid Khan and Dr. Chris J. Janse (Leiden Malaria Research Group) for providing transgenic *Plasmodium berghei* lines.

## Supporting information

S1 Text. Methods for generation and characterization of monoclonal antibodies to *P. falciparum* TRAP

S2 Text. Supplemental figure legends.

S3 Fig s1. Antibody epitope mapping.

S4 Fig s2. Flow cytometry analysis of antibody binding to TRAP transfectants.

S5 Fig s3. Antibody staining of sporozoites.

S6 Fig s4. Antibody competition.

S7 Fig s5. Effect of antigen disulfide reduction on antibody reactivity.

S8 Table s1. Summary of antibodies to P. falciparum TRAP

